# A region of Drosophila SLBP distinct from the histone pre-mRNA binding and processing domains is essential for deposition of histone mRNA in the oocyte

**DOI:** 10.1101/2020.04.16.030577

**Authors:** Jennifer Potter-Birriel, Graydon B. Gonsalvez, William F. Marzluff

**Affiliations:** Curriculum in Genetics and Molecular Biology, University of North Carolina at Chapel Hill, Chapel Hill, NC; Interdisciplinary Program in Biological and Genome Sciences, University of North Carolina at Chapel Hill, Chapel Hill, NC; Department of Biochemistry and Biophysics, University of North Carolina at Chapel Hill, Chapel Hill, NC; Department of Cellular Biology and Anatomy, Medical College of Georgia, Augusta University, Augusta, GA

## Abstract

During *Drosophila* oogenesis, large amounts of histone mRNA and proteins are deposited in the developing oocyte. These are sufficient for the first 14 embryonic cell cycles and provide the developing embryo with sufficient histone proteins until the zygotic histone genes are activated. The maternally deposited histone mRNA is synthesized in stage 10b of oogenesis after completion of endoreduplication of the nurse cells. Histone mRNAs are the only cellular mRNAs that are not polyadenylated, ending instead in a conserved stemloop instead of a polyA tail. The Stem-loop binding protein (SLBP) binds the 3’ end of histone mRNA and is essential for both the biosynthesis and translation of histone mRNA. We report that a 10 aa region in SLBP, which is not required for processing in vitro, is essential for transcription of histone mRNA in the stage 10b oocyte. In stage 10b the Histone Locus Bodies (HLBs) produce histone mRNAs in the absence of phosphorylation of Mxc, normally required for histone gene expression in S-phase cells. Mutants expressing this SLBP develop normally, produce small amounts of polyadenylated histone mRNA throughout development, but little histone mRNA in stage 10b resulting in death of the embryos in the first hr of development.

## INTRODUCTION

Histone mRNAs are the only known eukaryotic cellular mRNAs that are not polyadenylated. They end instead in a conserved stemloop which is formed by cotranscriptional endonucleolytic cleavage. Stemloop binding protein (SLBP) binds to the stemloop and participates in histone pre-mRNA processing. SLBP remains with the processed histone mRNA, and accompanies it to the cytoplasm and is essential for histone mRNA translation. Thus SLBP is stoichiometrically required for accumulation of functional histone mRNP. In cycling cells, histone mRNAs are typically cell cycle regulated, being synthesized just prior to entry into S-phase and rapidly degraded at the end of S-phase.

One exception to cell cycle regulation of histone mRNA is in oogenesis and early embryogenesis in many species which do not activate transcription during the early cell cycles. *Drosophila* provide a large maternal store of histone proteins and histone mRNAs that are synthesized in the nurse cells at the end of oogenesis, sufficient for embryos to develop until the 14^th^ cell cycle even in the absence of zygotic histone genes (Gunesdogan et al., 2010). In *Drosophila*, histone mRNAs are synthesized and cell-cycle regulated in the nurse cells as they go through multiple cycles of endoreduplication. At the end of the last S-phase, the histone mRNAs are degraded. Histone gene transcription is then activated in the absence of DNA replication and the histone mRNAs are synthesized in the nurse cells and then dumped into the oocyte (Ambrosio and Schedl, 1985; Ruddell and Jacobs-Lorena, 1985).

During the early embryonic cell cycles in many organisms there are rapid cycles consisting of S-phase followed by mitosis, which in *Drosophila* are as short as 5 minutes. The maternal histone mRNAs are stable during these cell cycles. At cycle 14 in *Drosophila*, the first G2 phase occurs, and the histone mRNAs, including the maternal and any newly synthesized zygotic histone mRNAs are degraded. An initial challenge for all animals in early development is to have sufficient histone proteins in the egg to remodel the sperm chromatin and then provide histones to package the replicating DNA as the embryo carries out the initial cell division cycles prior to activation of the zygotic genome. Different organisms have solved this problem in different ways (Marzluff et al., 2008).

There are ∼100 copies of each of the five *Drosophila* melanogaster histone genes, organized in a tandem unit of a 5 kB unit containing 1 copy each of the four core histone genes and a histone H1 gene (Bongartz and Schloissnig, 2019; Lifton et al., 1978; McKay et al., 2015). Each of these histone genes contains a polyadenylation signal after the histone processing signal. In cells with histone mRNA processing inhibited as a result of knockdown of factors required for histone pre-mRNA processing, polyadenylated histone mRNAs are produced (Wagner et al., 2007). The same alteration of histone gene expression occurs in flies with mutations in the histone processing machinery (Burch et al., 2011; Godfrey et al., 2006; Godfrey et al., 2009; Sullivan et al., 2001). Cultured *Drosophila* cells can proliferate in the absence of histone processing factors due to the polyadenylated histone mRNAs. Flies with mutations in the histone processing machinery develop past cycle 14 by producing polyadenylated histone mRNA. Depending on the time of onset of production and the extent of polyadenylated histone mRNA production resulting from the mutations, flies can develop to different stages. Severe mutants in SLBP are lethal, and produce polyadenylated histone mRNA starting in cycle 15. In contrast, flies with null mutations in U7 snRNA, which perdures through the initial larval stages, develop into adult flies that are sterile, and only produce large amounts of polyadenylated histone mRNA starting in the third instar larva (Godfrey et al., 2006).

Here we report that a region in SLBP distant from the RNA binding and processing domain is required for efficient processing of histone mRNA in vivo. Flies expressing this allele produce small amounts of polyadenylated histone mRNA throughout development. The animals are viable but maternal effect lethal due to the failure to express enough histone mRNAs at the end of oogenesis to deposit sufficient histone proteins and mRNAs into the egg.

## RESULTS

A schematic of the SLBP gene is shown in figure 1A, and a schematic of the domains of the SLBP protein are shown in Fig. 1B. The RNA binding and processing domains are sufficient for processing *in vitro*, while the N-terminal domain contains sequences required for translation of histone mRNA, import of the protein into the nucleus, and stability of the SLBP.

**Figure 1.**
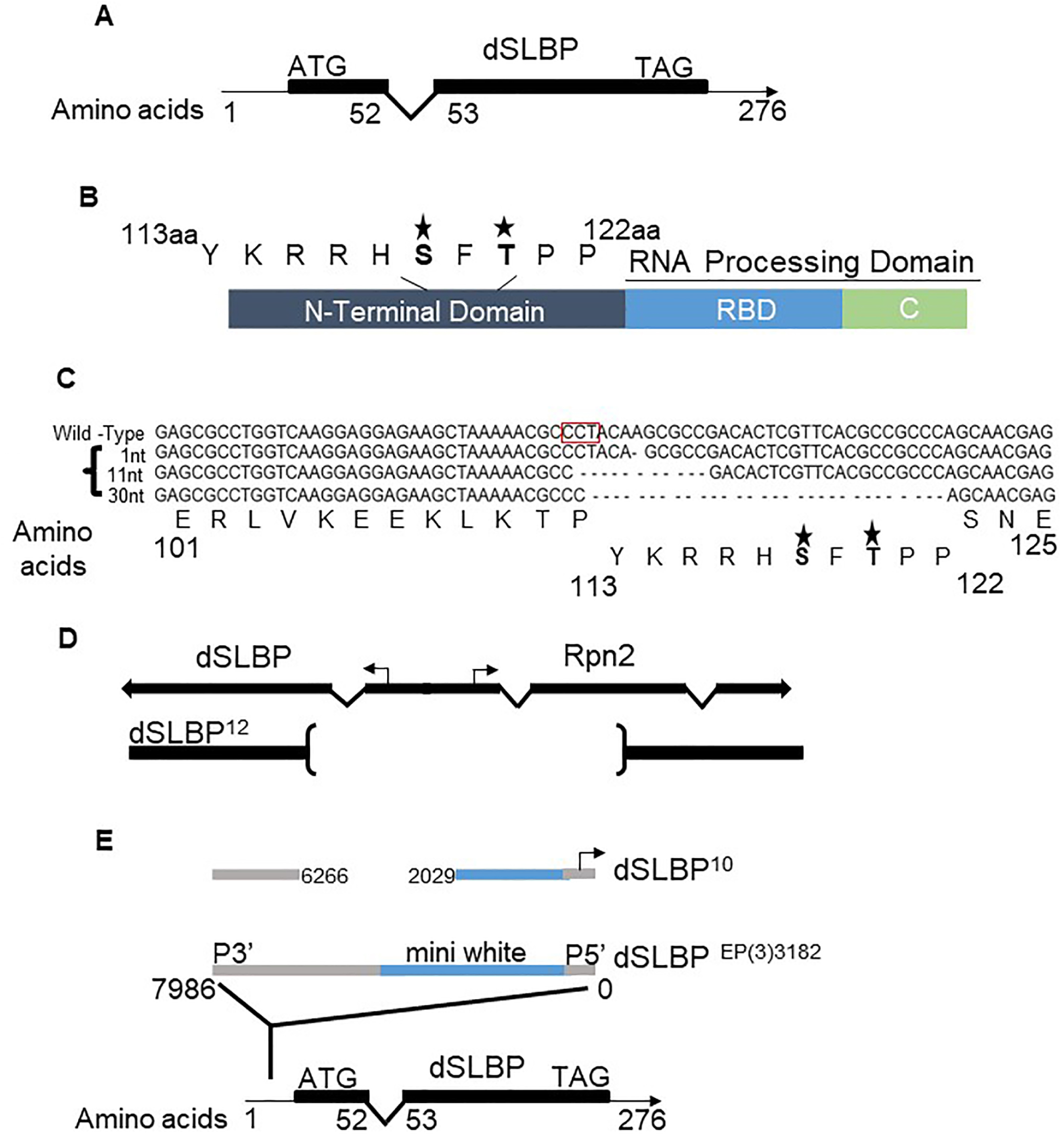
Schematic of SLBP structural domains, CRIPSR mutants and excision alleles. **A**. *Drosophila* SLBP contains two exons (black boxes) and one intron. **B**. Domain structure of SLBP showing the RNA processing domain composed of the RNA binding domain and the C-terminus. The N-terminal domain contains two known phosphorylation sites in the SFTPP motif located between amino acids 113 to 122. **C**. Sequence alignment of SLBP gene for each CRISPR deletion (1nt, 11nt and 30nt). Note that the SFTPP motif is deleted in the 30nt (10aa) deletion. The stars indicate the phosphorylated amino acids within the sequence. **D**. Diagram of the SLBP^12^ allele which contains a deletion extending from the coding region that includes the bidirectional promoter and a portion of the adjacent Rpn2 gene (Sullivan et al., 2001). **E**. Diagram of the SLBP^10^ allele which contains an insertion of part of a p element in the 5’ UTR region of SLBP (Sullivan et al., 2001). It is expressed from a cryptic promoter located near the P5’ end of the p element (Lafave and Sekelsky, 2011)

### Mutants in the SLBP gene

To study the biological function of SLBP, we used FLY-CRISPR Cas9 and a guide RNA with a PAM site at aa 112 to create a null mutant of SLBP in *Drosophila melanogaster*. We obtained two frameshift mutations resulting from a 1nt and an 11nt deletion (Fig. 1C). We also obtained a 30nt deletion which deleted amino acids 113 to 122 (Fig.1C). Interestingly, the 30nt (10aa) deletion removes two phosphorylation sites, one previously characterized by our group (Lanzotti et al., 2004) and the other, a site phosphorylated by Chk2, identified by Iampietro C., et al., 2014 (Fig. 1C). We also used two excision alleles previously characterized by our group, SLBP^12^, a molecular null that removes a portion of the coding region of SLBP and the adjacent Rpn2 gene (Fig. 1D), and SLBP^10^, a hypomorph which contains a fragment of the P element in the 5’UTR of SLBP (Fig. 1E). SLBP^10^ is a maternal effect lethal due to failure to produce enough histone mRNA at the end of oogenesis (Sullivan et al., 2001).

### Phenotype of an SLBP null mutant

SLBP is an essential factor for histone mRNA biogenesis and, not surprisingly, SLBP is essential for viability. Null mutants of SLBP in *C*.*elegans* die very early in development due to failure to produce zygotic histone mRNA (Kodama et al., 2002; Pettitt et al., 2002). In *Drosophila*, cells with greatly reduced levels of SLBP produce polyadenylated histone mRNAs since there is a cryptic polyadenylation site(s) 3’ of the SL in all *Drosophila* histone genes, which is utilized when histone pre-mRNA processing is blocked (Sullivan et al., 2001). There are large stores of maternal histone mRNA and protein stored in the egg, which are sufficient to support development through cycle 14 even in animals with the histone genes deleted, and in flies with histone mRNA processing factors mutated, including SLBP, development proceeds further due to the production of polyadenylated histone mRNA (Godfrey et al., 2006; Godfrey et al., 2009; Sullivan et al., 2001).

The original SLBP mutant we analyzed had an intact SLBP gene with a p element inserted in the 5’ UTR. This mutant, SLBP^15^, retained most of the p-element in the 5’ UTR, and died in the third instar larva, producing primarily polyadenylated histone mRNA after early embryogenesis (Godfrey et al., 2006; Sullivan et al., 2001). However SLBP^15^ is likely a hypomorph since there is a cryptic promoter at the end of the p element (Lafave and Sekelsky, 2011). We determined that the null mutants SLBP^Δ11^ and SLBP^Δ1^ died as first instar larvae when they were present with a deficiency of SLBP, SLBP^12^ (Fig. 2A), which removes the 5’ ends of both the SLBP gene and the adjacent Rrpn2 gene (Sullivan et al., 2001)(Fig. 1D). SLBP^Δ11^/TM3 Ser P[act-GFP] females were crossed with SLBP^12^/TM3 Ser P[act-GFP] males (Fig. 2A), and the reciprocal cross was also done. All the embryos with the mother contributing the SLBP^12^ allele hatched, but died very rapidly after hatching. In contrast when the male contributed the SLBP^12^ allele, the first instar larvae survived for several hours after hatching.

**Figure 2.**
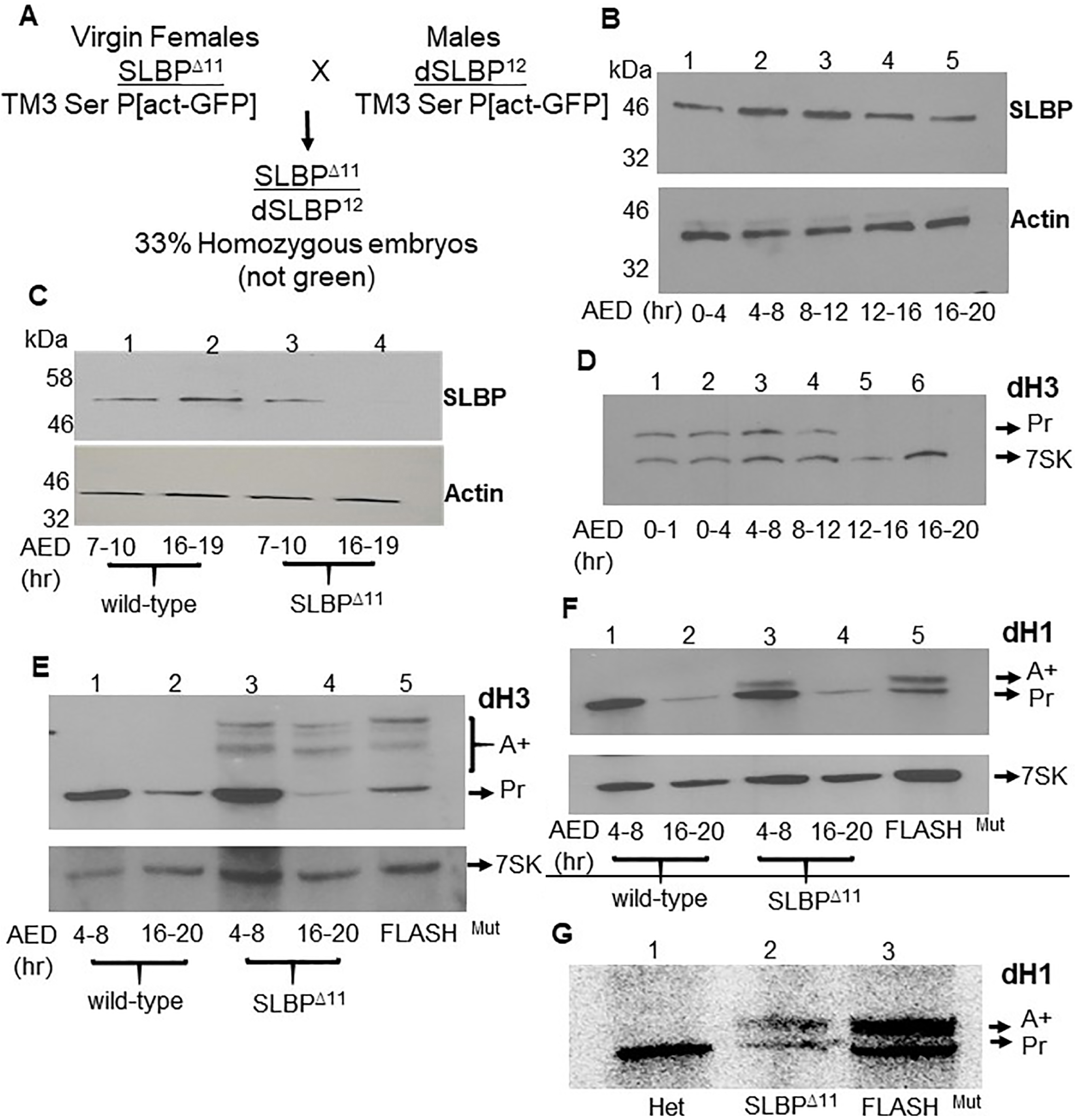
The null mutant (11nt deletion) dies in first instar larval stage, and produces polyadenylated histone mRNA starting early in development. **A**. SLBP^Δ11^/TM3 Ser P[act-GFP] females were mated with SLBP^12^/TM3 Ser P[act-GFP] males, and then mutant embryos not expressing GFP were selected starting 4 hrs deposition when GFP was expressed. Protein and RNA were extracted from either wild-type (w^1118^) or SLBP^Δ11^/SLBP^12^ embryos and SLBP detected by Western blotting and histone mRNA by Northern blotting. **B**. Western blot analysis of wild type (w^1118^) embryos collected every 4 hours of embryogenesis. Substantial amounts of SLBP are deposited in the egg [this will add this lane]. SLBP expression increases between 4-12 hr after embryo deposition (AED) and then declines. **C**. Western blot analysis of sibling wild-type embryos (lane 1-2) and null mutant embryos (lanes 3-4) collected between 7-10 hrs, and 16-19 hrs after AED. There are much lower levels of SLBP in the mutant embryos by 16-19 AED as the maternal supple is depleted. **D**. Expression of histone mRNA in embryogenesis. Total RNA from equal numbers of wild-type embryos was analysed by Northern blotting for histone H3 mRNA, with 7SK RNA analyzed as a loading control. Large amounts of histone mRNA is deposited in the egg (lane 1), levels increase at between 4-8 hrs AED and decrease later in embryogenesis. **E and F**. RNA was prepared from sibling wild-type (lane 1-2) and null mutant embryos 4-8 hrs and 16-20 hrs AED and analyzed by Northern blotting for histone H3 mRNA (E) and histone H1 mRNA (F). Lane 5 is analysis of a FLASH mutant embryo that expresses both processed and polyadenylated mRNA (Tatomer et al., 2016). Polyadenylated histone H3 and H1 mRNAs are detectable between 4-8 hrs of development. **G**. and are a large embryo Northern blot analysis demonstrate that homozygous embryos accumulate polyadenylated transcripts beginning at 4-8hr AED. **G**. Heterozygous and homozygous mutant 1^st^ instar larvae were collected and total RNA analyzed by Northern blotting for histone H1 mRNA. The filter was analyzed using a PhosphorImager.

We carried out molecular analyses with SLBP^Δ11^. We determined how long the maternal supply of SLBP protein lasted. Previously we did not have an antibody that was capable of detecting SLBP by Western blotting. We developed an affinity-purified antibody raised against full-length SLBP, which gave a single band in a Western blot. In normal embryos there are large amounts of SLBP in the egg, as previously reported by Lecuyer’s group (Iampietro et al., 2014). SLBP levels increased after activation of the zygotic genome and remained high during the period of active cellular proliferation from 4 to 12hrs after egg deposition (AED) (Fig.2B), after which it declined slowly until hatching. We collected embryos from WT and SLBP null mutant embryos from 7-10 hrs and from 16-19 hrs (AED) embryos. Western blot analysis of the collected embryos shows that while maternal SLBP declined during embryogenesis, in the null mutant SLBP was almost undetectable by 16-19 hrs.

We also analyzed *Drosophila* Histone H3 and Histone H1 (dH3 and dH1) mRNAs by Northern blotting. During early embryogenesis (up to 8 hrs) histone mRNAs are very abundant, but decrease by 12 hrs, and are very low after 16 hrs as DNA replication slows later in development (Fig. 2D). In SLBP^Δ11^ mutant embryos aged for 4-8 hrs there was some polyadenylated dH1 and dH3 mRNAs, indicating that histone pre-mRNA processing was inefficient starting early in embryogenesis, once all the histone mRNAs are zygotically expressed (Fig.2E-F). By 16hr of embryogenesis, the great majority of the dH3 mRNAs were polyadenylated (Fig. 2E, lane 4), although the total amount of H3 mRNA was much less than in early embryogenesis. Polyadenylated histone dH1 mRNA was present at similar amounts as histone H3 mRNA at 4-8 hrs, but there was a smaller amount of polyadenylated histone H1 mRNA later in embryogenesis.

The presence of polyadenylated histone mRNAs early in embryogenesis suggest that the mutant embryos didn’t have enough active SLBP to efficiently process the histone pre-mRNAs by 4-8 hrs of embryogenesis. We also collected homozygous and heterozygous first instar larvae to ask if they had any detectable polyadenylated transcripts. The northern blot was probed with dH1, and it shows that homozygous larvae contain both polyadenylated and processed histone mRNAs (Fig.2G). Note that the levels of SLBP decline much more slowly than histone mRNA during early embryogenesis, accounting for the perdurance of processed histone mRNA in the early larval stage.

### The 30nt (10aa) deletion is maternal effect lethal

Genetic analyses were done to ask whether the 30nt (10aa) deletion (SLBP^Δ30^) was viable over the deficiency of SLBP, SLBP^12^. The results showed that flies containing one allele of SLBP^Δ30^ expressing the 30nt (10aa) deletion were viable but maternal effect lethal. Previous work by our lab had shown that a hypomorphic allele of SLBP, SLBP^10^, had a similar maternal effect lethal phenotype, due to the failure to provide sufficient maternal supply of histone mRNA or protein in the egg. The embryos from SLBP^Δ30^ females showed a similar phenotype to SLBP^10^. None of these embryos developed past the syncytial stage and they died as a result of mitotic defects in the syncytial cycles, a similar phenotype to the SLBP^10^ embryos (Sullivan et al., 2001)(Fig. S1). The SLBP^Δ30^ embryos died within the first hr of development, while the SLBP^10^ embryos survived for 90 minutes.

The SLBP^10^ mutant has an insertion from the initial p element remaining in the 5’ UTR (Fig. 1E). Since we did not have an SLBP antibody when this mutant was first described we could not determine the levels of SLBP in this mutant. The SLBP^Δ30^ protein has a 10 aa deletion, between the N-terminus and the RNA binding domain (Fig.3A). The resulting protein has normal activity in histone pre-mRNA processing in vitro. The 10 aa deletion contains two phosphorylation sites, a threonine of unknown function which is phosphorylated on about 30% of the molecules (Borchers et al., 2006) and a serine, that is a Chk2 substrate (Iampietro C., et al., 2014). The Chk2 phosphorylated form plays a role in destruction of SLBP allowing the identification of defective nuclei during the syncytial stages of development.

**Figure 3.**
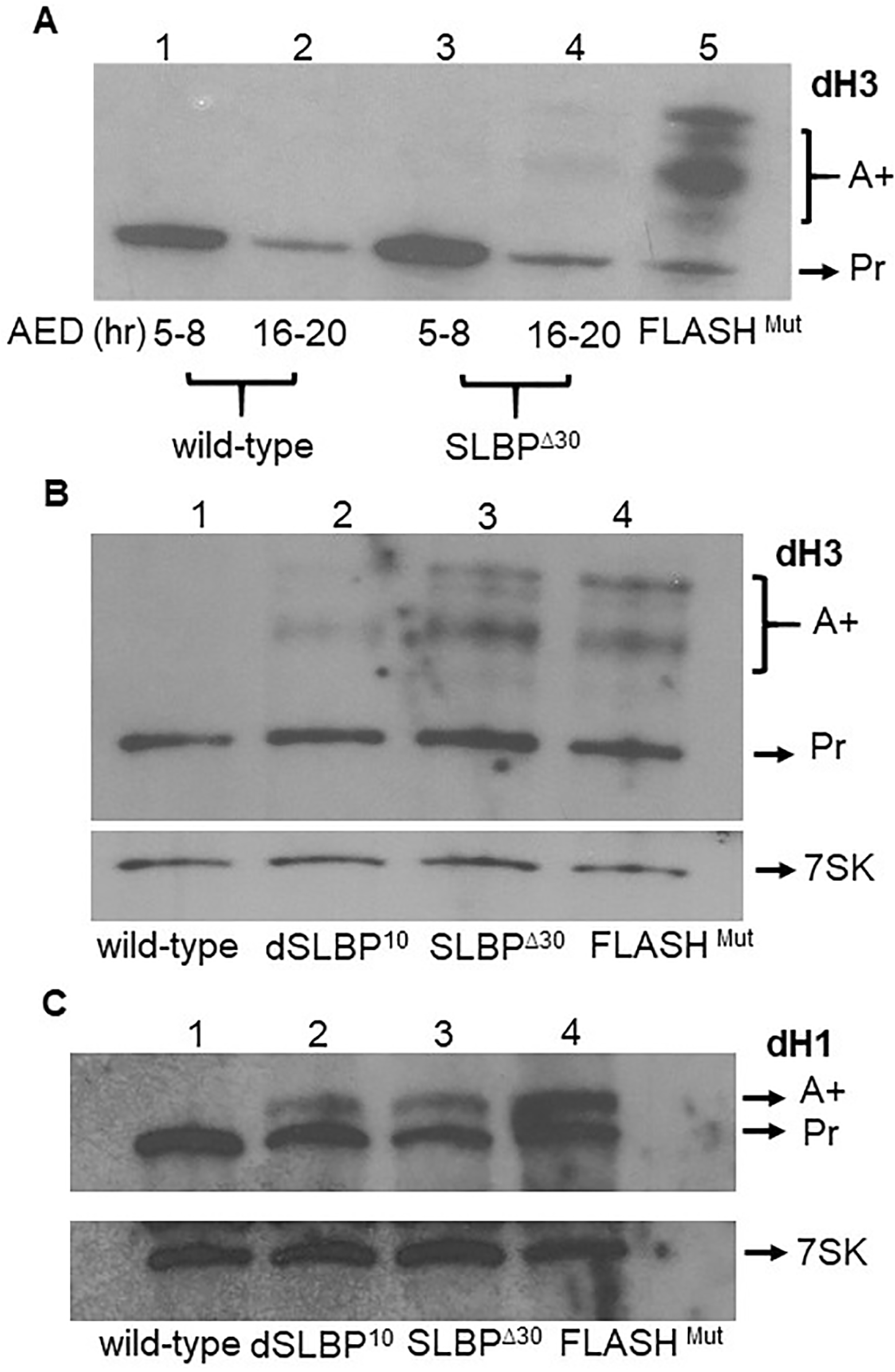
A fraction of Histone mRNAs in SLBP^Δ30^ are polyadenylated throughout *Drosophila* development. **A**. Wild type (w^1118^) and homozygous embryos SLBP^Δ30^ /SLBP^12^ were collected from 5-8 and 16-20hrs (AED). SLBP^Δ30^ begins to accumulate polyadenylated dH3 at low levels in 5-8 embryos, and it is a substantial fraction of the histone H3 mRNA at 16-20hrs (AED) Actin was assayed as a control. **B. and C**. 3rd instar larvae were collected from w^1118^ and homozygous (GFP-) mutants SLBP^Δ30^/SLBP^12^ and SLBP^10^/SLBP^Δ11^. Equal amounts of RNA were probed was probed for dH3 (B) and dH1 (C) together with 7SK RNA as a loading control. Lane 4 is the FLASH mutant as a positive control.

### Expression of polyadenylated histone mRNA throughout development in SLBP^Δ30^

We examined whether the hypomorphic mutants also produce polyadenylated histone mRNAs at other developmental stages. We prepared total RNA from different times in embryogenesis, and third instar larvae and analyzed it by Northern blotting. Polyadenylated histone mRNAs were detected in the SLBP^Δ30^ mutant by the end of embryogenesis (Fig.3A, lane 4), and continued to be expressed throughout the larval stages (Fig.4B-4C) and in ovaries (Fig.3D). The SLBP^10^ mutant embryos expressed a lower amount of polyadenylated histone mRNA than did the SLBP^Δ30^ mutant, and in 3^rd^ instar larvae.

**Figure 4.**
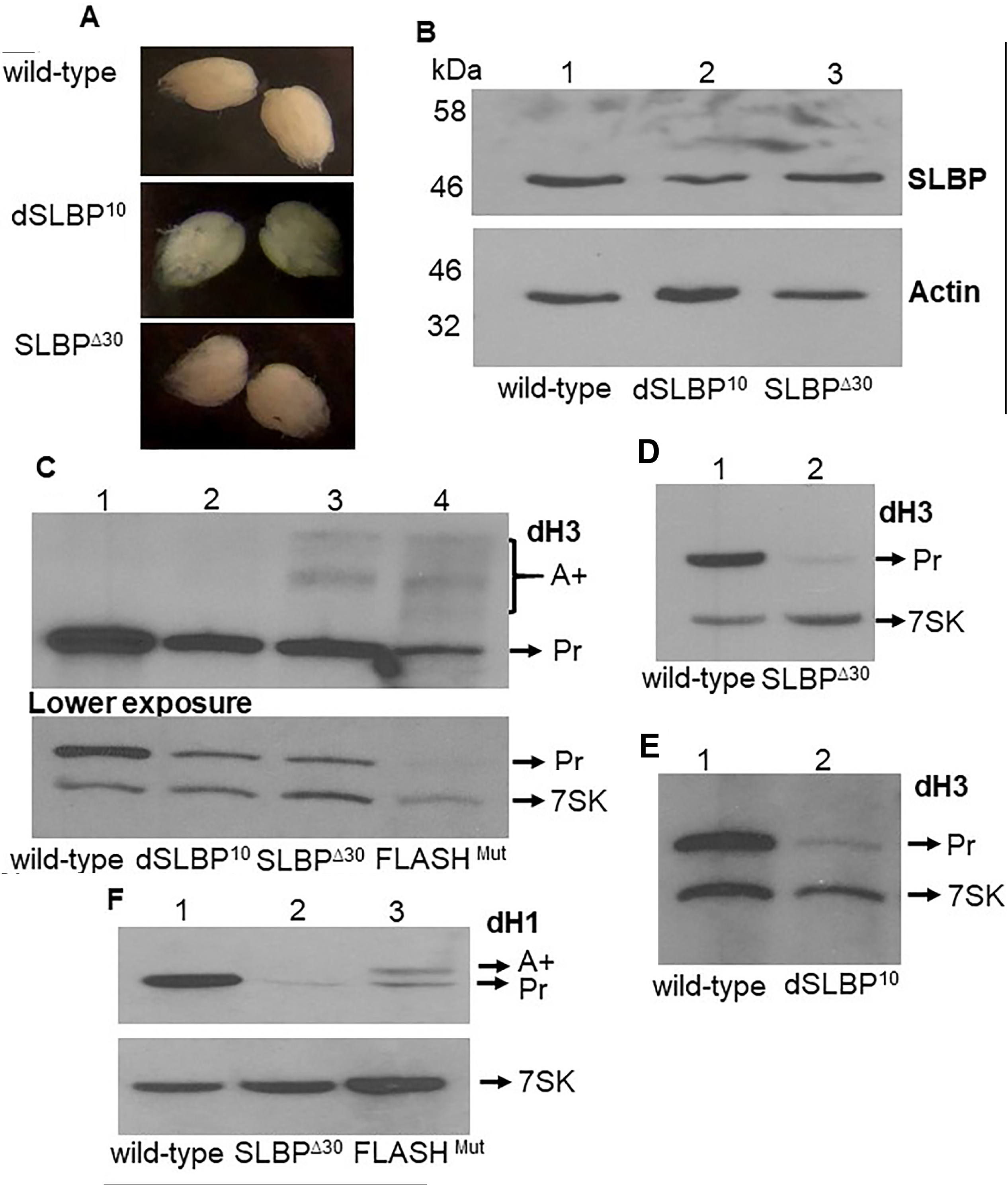
SLBP and histone mRNAs in ovaries from hypomorphic SLBP^10^ and SLBP^Δ30^ flies. **A**. Females were aged for 4-6 days, the dissected ovaries from both SLBP^10^ and SLBP^Δ30^ hypomorphs. They are similar in size to those from wild-type flies. **B**. Ovaries from SLBP^10^ and SLBP^Δ30^ homozygous females and wildtype flies and analyzed for SLBP protein by western blotting using actin as a loading control. **C**. Histone H3 mRNA from ovaries from SLBP^Δ30^ and SLBP^10^ was analyzed by Northern blotting for H3 mRNA using 7SK RNA as a loading control. RNA from the FLASH mutant was used as a positive control. **D. and E**. Northern Blot analysis showed reduced levels of dH3 mRNA in embryos collected from 30 to 45 minutes (AED) from SLBP^Δ30^ (D) or SLBP^10^ (E) females. 7SK RNA was probed as a control. **F**. dH1 mRNA was analyzed from the same RNA sample from SLBP^Δ30^ embryos with FLASH mutant as a control. During this time the embryo only contains the maternal supply loaded into the egg.

### Effect of hypomorphic mutants on ovary function

The ovaries of SLBP^10^ and SLBP^Δ30^ females were normal in appearance, and similar in size to wild-type ovaries (Fig.4B), and the animals laid normal numbers of eggs. The ovaries from the mutants had normal looking germaria, and the different stages of egg chamber maturation were similar in both the mutant and wild-type animals. We examined the SLBP protein levels in both SLBP^Δ30^ and SLBP^10^ ovaries by Western blotting and histone mRNA by Northern blotting. Females expressing only SLBP^10^ produced a lower amount of SLBP protein compared to wild type (Fig.4C, lane 2). In contrast, SLBP^Δ30^ ovaries contained normal levels of SLBP (Fig.4C, lane 3). A northern blot analysis showed reduced levels of dH3 mRNA in ovaries from both mutants. The SLBP^Δ30^ ovaries expressed some polyadenylated dH3 mRNA as well as properly processed mRNA, but the SLBP^10^ mutant ovaries expressed much less polyadenylated mRNA than the SLBP^Δ30^ ovaries (Fig.4D).

### SLBP^Δ30^ females deposit very low levels of histone mRNAs into eggs

It is known that histone mRNAs (Ambrosio and Schedl, 1985; Ruddell and Jacobs-Lorena, 1985) and proteins are synthesized at the end of oogenesis and loaded into the egg. These proteins and mRNAs are utilized during the syncytial stages of embryogenesis prior to activation of the zygotic genome, since embryos lacking any histone genes develop through cycle 14 (Gunesdogan et al., 2010). We previously showed that eggs from SLBP^10^ mutant ovaries contain about 10% as much histone mRNA (Lanzotti et al., 2002) and likely protein as wild type embryos. We determined if the maternal supply of histone mRNA was affected in embryos laid by females expressing SLBP^Δ30^ by Northern blotting. Embryos from 30 to 45 minutes AED were collected for this analysis and the amount of histone H3 and Histone H1 mRNA deposited into the egg were determined (Fig. 4E and F). Only very small amounts of each RNA (<5% of WT) were detected in the eggs from SLBP^Δ30^ embryos, less than was present in the SLBP^10^ eggs (Fig.4G), consistent with these embryos dying earlier than the SLBP^10^ embryos.

### Expression of histone mRNAs during different stage of oogenesis in the hypomorphs

Development of the oocyte occurs within a germarium in the ovary. A germ-line stem cell undergoes 4 cycles of division resulting in 16 cells. One of these cells becomes the oocyte, while the other 15 cells become nurse cells. The nurse cells undergo multiple rounds of endoreplication. The nurse cells and oocyte are surrounded by a layer of somatic follicle cells whose numbers dramatically increase during egg chamber maturation. Thus, Western and Northern blots of the whole ovary detect the mRNAs and proteins in both the oocyte, nurse cells and follicle cells.

To study histone mRNA metabolism in the developing egg chamber, we performed single molecule fluorescent in situ hybridization (smFISH) using Stellaris probes against the coding region of histone H3 (H3-coding). Consistent with previous results, in the germanium and early stage egg chambers, H3-coding in situ signal could be observed in some but not all cells (Fig.S2A, B). There was cytoplasmic mRNA in some cells, and these cells also had labeled foci in the nuclei. Early-stage egg chambers in both the SLBP^10^ (Fig. S2C) and SLBP^Δ30^ (Fig. S2D) mutants developed normally and was similar to the wild-type in having both cytoplasmic histone mRNA and nuclear foci in some cells. Co-staining with FLASH, a component of the HLB and the histone coding probe, showed that the nuclear foci were HLBs (Fig. S2C’,D’). In order to confirm that these cells correspond to those that are in S-phase, egg chambers were stained with an antibody against FLASH, and a phospho-Mpm2 antibody to identify S-phase cells (White et al., 2007). Although FLASH positive nuclear foci could be detected in all cells, only a fraction of these nuclear foci were positive for phospho-Mpm2 in wild-type cells (Fig.5A, A’, A’’). The phospho-Mpm2 positive cells correspond to follicle cells and endoreplicating nurse cells that are in S phase.

**Figure 5.**
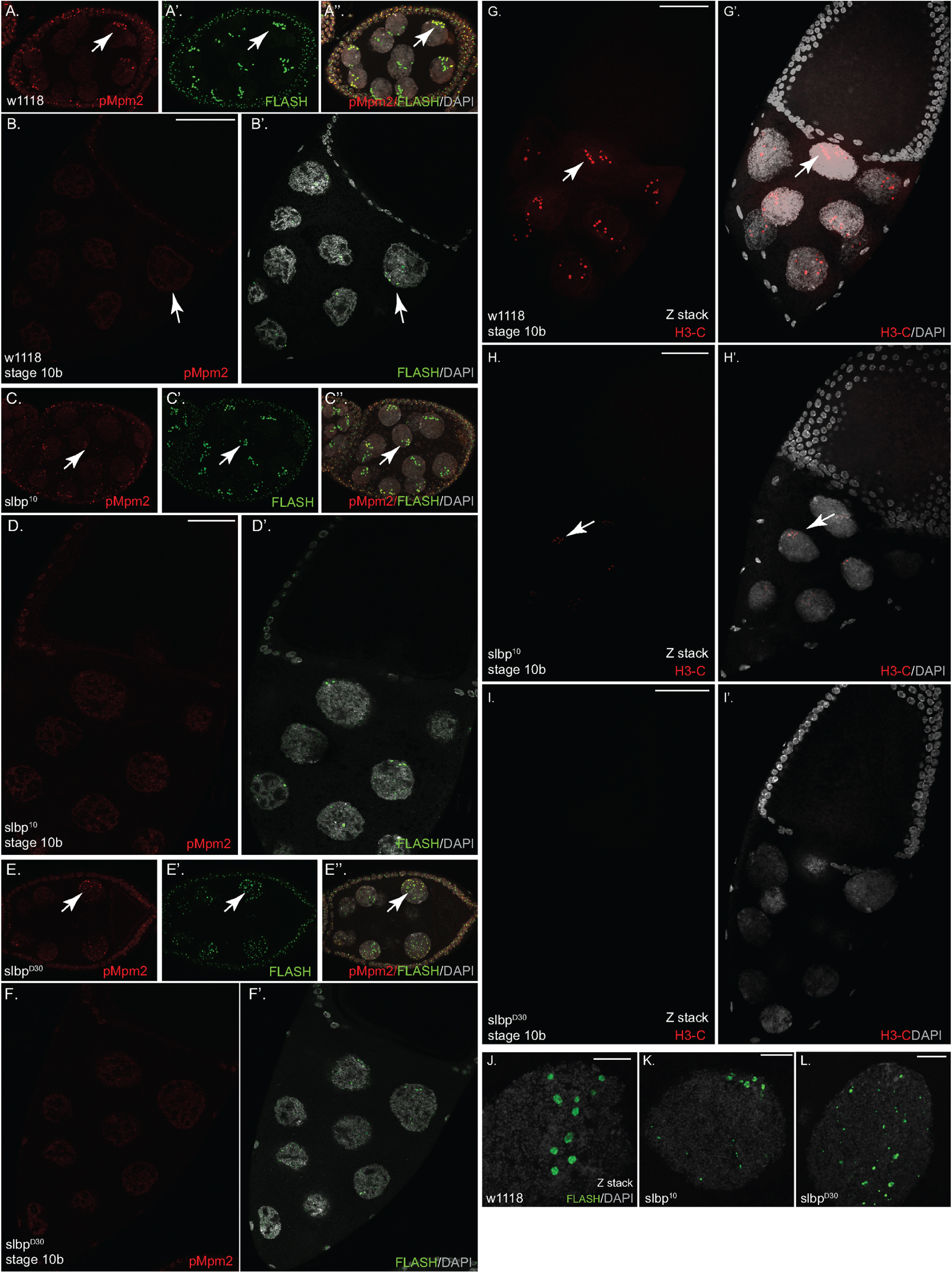
Histone expression in defective in SLBP^10^ and SLBP^Δ30^ stage10b egg chambers. **A**. An early stage wild-type egg chamber stained with the Mpm2 (A, red), and FLASH (A’, green) antibodies. A merged image along with DAPI (grey) is shown in A’’. Arrows indicate an mpm2 positive HLB. **B**. A wild-type stage 10b egg chamber stained with Mpm2 (B, red) and FLASH (B’, green) is shown. The arrow indicates nuclear FLASH foci. **C**. An early stage SLBP^10^ egg chamber stained with the Mpm2 (C) and FLASH (C’) antibodies is shown. A merged image along with DAPI is shown in C’’. Arrows indicate an mpm2 positive HLB. **D**. A SLBP^10^ stage 10b egg chamber stained with Mpm2 (D) and FLASH (D’) is shown. **E**. An early stage SLBP^Δ30^ egg chamber stained with the Mpm2 (E) and FLASH (E’) antibodies is shown. A merged image along with DAPI is shown in E’’. Arrows indicate an mpm2 positive HLB. **F**. A SLBP^Δ30^ stage 10b egg chamber stained with phospho Mpm2 (F) and FLASH (F’) is shown. **G**.**-I**. Stage 10b egg chambers from wild-type (G), SLBP^10^ (H) or SLBP^Δ30^ (I) processed for in situ hybridization using probes against the H3 coding region are shown (H3-C, red). The images represent a Z stack. A merged image with DAPI is shown in G’, H’ and I’. Arrows indicate nuclear H3 foci present in all nuclei in wild-type 10b egg chambers. There are a small number of nuclei in SLBP^10^ nurse cell nuclei which have weak foci (arrow). **J**.**-L**. A Z stack of a single nurse cell nucleus from wild-type (J), SLBP^10^ (K) or SLBP^Δ30^ (L) stained with the FLASH antibody (green) and counter-stained with DAPI (grey) is shown.

### Histone mRNAs are expressed from mpm-2 negative HLBs for deposition into the egg

Endoreplication of nurse cells is complete by stage 10. FLASH positive nuclear foci (HLBs) could be detected in nurse cells of stage 10b egg chambers, but most of these (86%) were negative for phospho-Mpm2 (Fig. 5B, B’). In the 14% of wild-type stage 10b oocytes that were positive for mpm2, all the HLB were positive (Fig. S2E, E’, E”). Despite not being in S-phase, these cells displayed abundant H3-coding in situ signal (Fig.5G,G’) at nuclear foci in all cells. Thus, at stage 10b, histone gene expression is turned on in all nurse cells in the absence of DNA replication, and in the great majority of the egg chambers the HLBs are mpm2 negative.

We next compared histone expression between wild-type and SLBP mutant egg chambers. Histone mRNA expression in the germarium and early stage egg chambers was similar between wild-type, SLBP^10^, and SLBP^Δ30^ mutants (Fig. S2 C, D). In stage 10b egg chambers and later, the mutants differed markedly from the wild-type. Although abundant histone transcription foci could be detected in the nurse cells of wild-type egg chambers, transcription foci were greatly reduced (SLBP^10^) or not detectable (SLBP^Δ30^) in the SLBP mutants. The SLBP^10^ mutant displayed a weaker phenotype and in these egg chambers some transcription foci could be detected (Fig. 5H,H’). However, the level of H3-coding in situ signal at the nuclear foci was significantly reduced in comparison to wild-type (Fig. 5H, H’), and the cytoplasmic signal was also much weaker than wild-type. SLBP^Δ30^ mutants displayed a more severe phenotype and the in situ signal was not detectable in most stage 10b egg chambers (Fig.5I, I’). The HLBs also differed in morphology in the stage 10B nurse cells in the wild type and the two mutants. The FLASH foci were larger and there were fewer of them in the WT stage 10B egg chambers, while in the two mutants, the foci were smaller and there were many more per cell in both mutants, with the SLBP^Δ30^ being more severely affected than the SLBP^10^ mutants (Fig. 5J-L).

Nurse cell dumping in late stage egg chambers results in deposition of histone mRNA into the oocyte. Consistent with the lack of histone expression observed in stage 10b egg chambers, and the amount of maternal histone mRNA present in the early embryos, both mutants contained substantially reduced histone mRNA in the oocyte of late-stage egg chamber (Fig.6C, F, I). Slightly more histone mRNA signal could be detected in the oocyte of late-stage SLBP^10^ mutants in comparison to SLBP^Δ30^ mutants (Fig. 6F, I).

**Figure 6.**
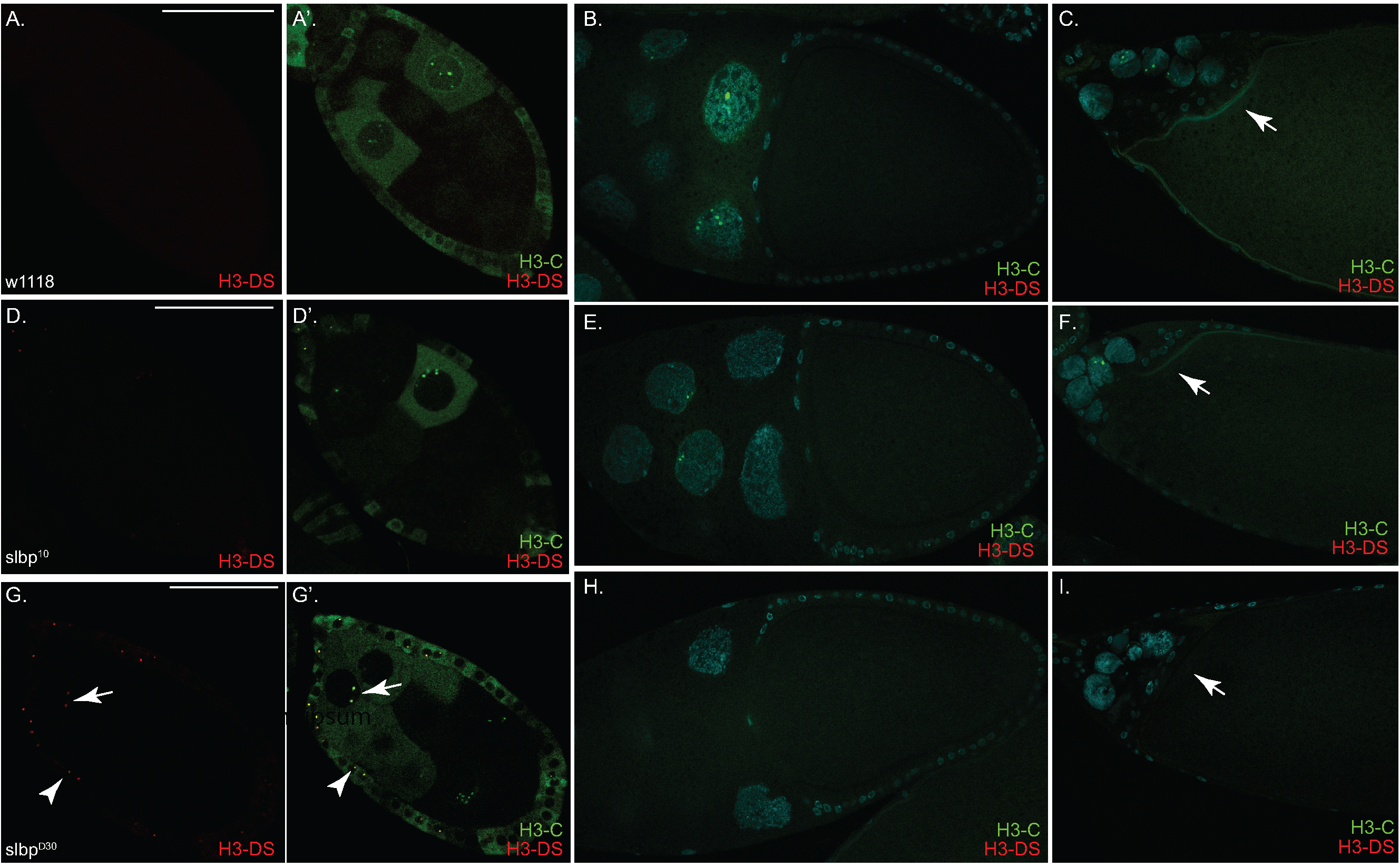
Misprocessed histone transcripts can be detected in early-stage SLBP^Δ30^ egg chambers. Egg chambers from wild-type **(A-C)**, SLBP^10^ **(D-F)** or SLBP^Δ30^ **(G-I)** mutant ovaries were processed for in situ hybridization using probes against the coding region of H3 mRNA (H3-C, green) and against a region downstream of the coding region (H3-DS, red). H3-DS signal was mostly detected in early stage SLBP^Δ30^ egg chambers. The H3-DS signal was most often seen in follicle cells (G, arrow head) but could occasionally also be observed in the nuclei of SLBP^Δ30^ mutant egg chambers (G, arrow).

We next determined whether histone mRNA processing was disrupted in the germarium of SLBP mutants. Egg chambers were processed with two probes, the H3-coding probe (H3-C) and a probe that hybridizes to a region downstream of the coding region (H3-DS). This region is not expressed when nascent histone transcripts are correctly processed, but is expressed in mutants where processing is defective or occurs with slower kinetics (Tatomer et al., 2016). In early stage wild-type and SLBP^10^ mutant egg chambers, H3-C signal was observed in nuclear foci and also diffusely in the cytoplasm of nurse cells and follicle cells in S-phase. Minimal signal was detected with the H3-DS probe in the wild type or SLBP^10^ egg chambers (Fig.6A, A’, D, D’). By contrast, in SLBP^Δ30^ mutants, H3-downstream probe signal was observed in some of these egg chambers. Often signal from the downstream probe was detected in follicle cell nuclei (Fig.6G, G’, arrowhead). Occasionally signal from the downstream probe could also be detected within nurse cell nuclei (Fig.6G, G’, arrow). At stage 10b, H3-coding signal, but not H3-downstream signal, was detected in wild-type egg chambers (Fig. 6B), both as transcription foci in nuclei and in the cytoplasm near the oocyte. At the same stage, both mutants displayed reduced H3-coding signal, both in nuclear foci and the cytoplasm. Signal from H3-DS was also minimal in both mutants (Fig.6E,H), consistent with an overall reduction in transcription. As nurse cell dumping continues, the nurse cells continue to have transcription foci and large amounts of histone mRNA accumulates in the wildtype oocyte (Fig. 6C), but much lower amounts of histone mRNA accumulates in the oocyte in the SLBP^10^ mutant and still lower amounts in the SLBP^Δ30^ mutant.

### Mislocalization of SLBP in SLBP^Δ30^ mutants

Western blot analysis indicates that SLBP protein is expressed in ovaries from SLBP^10^ and SLBP^Δ30^ mutants (Fig. 3C). In order to more precisely characterize the molecular nature of the defect, we examined the localization of SLBP in wild-type and mutant ovaries by immunofluorescence. SLBP could be detected in cytoplasmic and nuclear compartments in wild-type egg chambers, SLBP was present in significantly higher concentration in the nuclei of both nurse cells and follicle cells during all early stages of oogenesis as well as stage 10B (Fig. 7A). Note that SLBP is present in all nurse cells suggesting that its amounts do not change during the cell cycle in these cells (Fig.7A). In the SLBP^10^ mutant, the level of SLBP protein was reduced in early-stage and stage 10b egg chambers, although its localization was normal and the nuclear enrichment of SLBP was still evident in the SLBP^10^ mutant background (Fig.7 B, E). In contrast, although the level of SLBP^10^ was similar to the wild-type, the nuclear enrichment of SLBP in nurse cells and follicle cells was greatly reduced in SLBP^Δ30^ mutant egg chambers (Fig. 7C, F, G). Collectively, these results suggest that abundant nuclear SLBP in stage 10b egg chambers is required for expression of histone genes.

**Figure 7.**
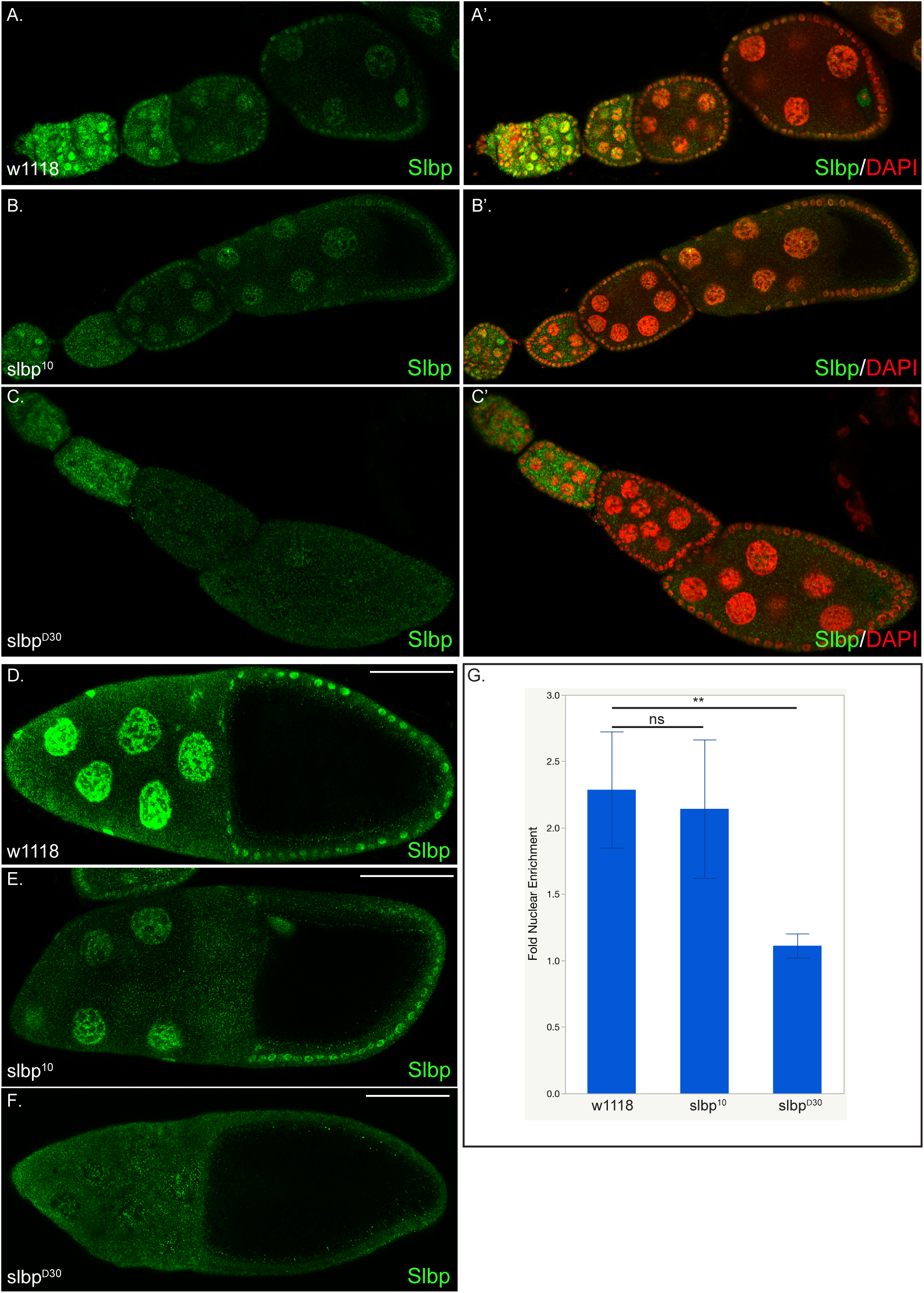
SLBP is mislocalized in SLBP^Δ30^ egg chambers. Egg chambers from wild-type **(A, A’, D)**, SLBP^10^ **(B, B’, E)** or **SLBP**^**Δ30**^ **(C, C’, F)** mutant ovaries were processed for immunofluorescence using an antibody against SLBP (green). The images shown in A’, B’ and C’ are counterstained with DAPI (red). In wild-type and SLBP^10^ mutants, Slbp protein is enriched within the nucleus (G). Note there is less SLBP present in SLBP^10^ egg chambers There is no enrichment of SLBP in the nuclei of SLBP^Δ30^ mutants in either the nurse cells or follicle cells (G). ns represents not significant. ** p <0.05, unpaired *t* test.

### Phosphorylation of the SFTTP region is not required for histone mRNA deposition

The 10 amino acid region that was deleted from SLBP contains the sequence SFTPP, which is similar to the sequence involved in regulating mRNA half-life in vertebrate SLBPs. We previously showed that the T is phosphorylated in SLBP expressed in baculovirus expressed SLBP and in cultured cells (Torres et al., 2005), and Lecuyer and coworkers showed that the S is phosphorylated by Chk2 during syncytial stages of embryogenesis as part of a quality control mechanism that identifies defective nuclei (Iampietro et al., 2014). Previous we had shown that mutation of SFTPP to SFAPP, and this SLBP rescued an SLBP mutant and the females were fertile (Lanzotti et al., 2004). We constructed SLBP mutants where we changed the sequence to AFTPP and AFATP and these SLBP genes were integrated by recombinase mediated cassette exchange (RMCE) at the site 25C in the 2nd chromosome (Fig. S4A). These mutant SLBPs rescued the null mutations, restoring viability, and both males and females were fertile. There was also no polyadenylated histone mRNA detected in flies expressing only the mutant SLBPs (Fig. S4B). We conclude that the phosphorylation of the SFTPP region is not relevant to the deposition of maternal histone mRNA into the oocyte. We also found that mutations of the C-terminal phosphorylations, which are necessary for phosphorylation in vivo, were also not essential in vivo (fig. S4C-E).

## DISCUSSION

Histone mRNAs and proteins are expressed in large amounts during S-phase. Normally in *Drosophila*, processing of the 3’ end of histone mRNAs is very efficient, and in metazoans is tightly coupled with transcription termination (Eaton et al., 2018; Tatomer et al., 2016). The one time in the *Drosophila* life cycle that histone mRNAs are expressed at high levels in the absence of DNA replication, is at the end of oogenesis, when the histone mRNAs deposited into the egg are synthesized (Ambrosio and Schedl, 1985; Ruddell and Jacobs-Lorena, 1985). This may also be the time when demand for histone mRNA is highest in the *Drosophila* life cycle, as enough histone mRNA and histone protein are deposited into the egg to provide enough histones for the first 14 cell cycles. The mechanism by which the maternal stores of histone mRNA are synthesized outside of S-phase are not well understood.

SLBP is a critical factor for histone mRNA biogenesis and function, since it is required for histone pre-mRNA processing, and is stoichiometrically bound to cytoplasmic histone mRNA, where it is required for translation. Thus large amounts of SLBP are required not only for efficient processing but also for accumulation of large amounts of cytoplasmic histone mRNA. In vitro, processing requires only C-terminal 89 amino acids of SLBP, the RNA binding domain and the C-terminal 17 amino acids. While the requirements in SLBP for efficient histone pre-mRNA processing in vitro are well understood (Dominski et al., 2005; Dominski et al., 2002; Sabath et al., 2013; Skrajna et al., 2017; Skrajna et al., 2018), whether there are additional requirements in SLBP for efficient processing in vivo is not known. Here we show that a 10 aa region in the amino terminal of SLBP is essential for efficient histone 3’ end formation in vivo, and is required for deposition of maternal histone mRNA into the oocyte.

We had previously identified a hypomorph, SLBP^10^, which had a similar maternal lethal phenotype. It also produced some polyadenylated histone mRNA during development, and was female sterile, and only deposited small amounts of maternal histone mRNA in the egg (Lanzotti et al., 2002; Sullivan et al., 2001). This mutant produces lower amounts of wild-type SLBP protein (Fig. 3), although we did not have the reagents to determine that at the time we discovered it. The SLBP^10^ females deposit more histone mRNA into egg, and develop farther then the SLBP^Δ30^ embryos, although only have ∼10% the amount of wild-type animals (Lanzotti et al., 2002).

### How are maternal stores of histone mRNA produced independent of DNA replication?

We confirmed the initial reports of 35 years ago (Ambrosio and Schedl, 1985; Ruddell and Jacobs-Lorena, 1985) that maternal histone mRNAs were synthesized after the completion of nurse cell endoreduplication. At that time, the nurse cells start to “dump” large amounts of cytoplasmic contents (ribosomes, translation factors, maternal mRNAs) as well as localizing specific mRNAs into the developing oocyte. The other major component of the oocyte, yolk proteins, is synthesized by the follicle cells and endocytosed into the oocyte.

Although histone mRNAs are expressed by nurse cells that are in S-phase, prior to stage 10b, these mRNAs are not transported into the oocyte, but are degraded at the end of each S-phase. This is in contrast to mRNAs such as *bicoid, gurken* and *oskar* which are localized at specific regions within the developing oocyte and are critical to the development of the egg and embryo (Berleth et al., 1988; Ephrussi et al., 1991; Kim-Ha et al., 1991; Neuman-Silberberg and Schupbach, 1993). It is therefore likely that accumulation of histone mRNA within the oocyte requires nurse cell dumping, an actin-driven process that begins at stage 10b (Buszczak and Cooley, 2000). During dumping, nurse cell nuclei undergo apoptosis and their cytoplasmic contents are transferred into the egg. Interestingly, histone genes continue to be expressed at high levels in nurse cells even during late stages of egg chamber maturation. Thus, although some of the nurse cells are undergoing apoptosis, the ones that remain continue to transcribe histone genes. Although germ plasm mRNAs such as *oskar* and *nanos* continue to accumulate at the posterior of the oocyte during late stage of egg chamber maturation, this appears to rely on active localization pathways rather than continued transcription (Sinsimer et al., 2011; Snee et al., 2007).

The ovaries of both mutants are similar in size to wild-type ovaries and the structure of the germarium is normal. The development of the nurse cells is normal, with large amounts of histone mRNA in the cytoplasm in S-phase, with most of that mRNA properly processed. There is minimal histone mRNA (by in situ hybridization) at the start of Stage 10B of oogenesis. In wild type animals, histone mRNA starts to accumulate in the nurse cell cytoplasm and also in foci in the nuclei which represent the site of transcription of the histone genes at the histone locus body. Histone mRNA then starts to appear in the oocyte and rapidly accumulates throughout the oocyte (Fig. 6).

The HLB serves to concentrate factors required for both transcription and pre-mRNA processing at the histone genes. Histone gene transcription is normally activated after phosphorylation of the core HLB factor, Mxc, by Cyclin E/Cdk2 (White et al., 2007). In stage 10B oocytes, all the nurse cells nuclei produce histone mRNA in the absence of phosphorylation of Mxc. In wild-type egg chambers this mRNA is properly processed and deposited in the oocyte bound to SLBP.

### Effects of mutants which have defects in histone pre-mRNA processing

Mutations that affect histone pre-mRNA processing result in two distinct changes in histone gene expression we can detect by in situ hybridization. Normally histone mRNAs are rapidly processed and transcription terminated close to the processing signal. If histone pre-mRNA processing is slow, then transcription continues past the normal transcription termination site, allowing detection of the nuclear foci with a probe 3’ of the normal processing site. If those transcripts are then processed to produce polyadenylated mRNAs, then there is also polyadenylated histone mRNA detected in the cytoplasm. Alternatively the longer transcripts can ultimately by processed normally at the stemloop in which case little polyadenylated mRNA is procduced (Tatomer et al., 2016) or if they are never processed then they ultimately may be degraded in the nucleus. Defects in SLBP, due to low amounts of SLBP, results in production of polyadenylated histone mRNAs. The polyadenylated histone mRNAs produced in the SLBP^10^ and SLBP^Δ30^ mutants during development is likely a result of low concentrations of SLBP in the nucleus, slowing histone pre-mRNA processing,

One difference in both the maternal effect lethal mutants we found in stage 10b egg chambers was a low concentration of SLBP in the nucleus, due to inefficient nuclear localization in the SLBP^30^ mutant and low levels of total SLBP in the SLBP^10^ mutant. This lack of SLBP apparently leads to low levels of histone gene transcription and disruption of normal HLB structure in the stage 10B egg chambers where transcription occurs when Mxc is not phosphorylated. These results suggests that SLBP may be required for histone gene transcription from the HLB when there is not phosphorylation of Mxc.

The failure of SLBP^Δ30^ mutants to deposit histone mRNAs in the egg, may be a result of the mutant protein being predominantly in the cytoplasm. The region that is deleted is predicted to be part of a nuclear localization signal, and in contrast to mammalian SLBP, which has several potential nuclear localization signals (Erkmann et al., 2005), this is the only predicted canonical nuclear localization signal in *Drosophila* SLBP. Although histone mRNAs are not deposited in the egg, there is substantial amounts of SLBP in the eggs from the SLBP^10^ hypomorph (not shown). This is consistent with the finding that there is SLBP in the cytoplasm of the nurse cells even though mature histone mRNAs are not produced.

### Differences between requirements for histone pre-mRNA processing in vitro and in vivo

The SLBP^Δ30^ hypomorph is expressed at similar levels to wild-type SLBP, and processing is inefficient in vivo. We tested whether other SLBP mutants in phosphorylation sites, which are inefficient in processing in vitro, are also inefficient in vivo. The C-terminal 17 amino acids of SLBP are heavily phosphorylated in vivo, with four serines phosphorylated. In addition ser 270 in the conserved TPNK is phosphorylated in *Drosophila* and in mammals. Phosphorylation of the C-terminus is essential for processing in vitro, and phosphomimetic glutamic acid, only partially restores processing activity (Skrajna et al., 2017). However, when these proteins in vivo, and they both rescued the null mutant of SLBP, as effectively as the wild-type protein did (Fig. S4D,E). No polyadenylated histone mRNA was detected and both males and females were fertile.

Together these studies point emphasize that the effect of specific mutations in vitro on specific reactions may be very different from their effects in vivo where there are many more constraints on what must occur to carry out reactions efficiently at specific subcellular localizations.

## METHODS

### Fly stocks

w[1118] was used as the wild-type stock (Bloomington stock center; #5905). The shRNA strains were: *eb1* shRNA used as a control (Bloomington stock center; #36680, donor TRiP) and *slbp* shRNA (Bloomington stock center; #56876, donor TRiP). shRNA expression was driven using P(w[+mC]=matalpha4-GAL-VP16)V37 (Bloomington Stock Center, #7063; donor Andrea Brand).

### Antibodies

The following antibodies were used for immunofluorescence: rabbit anti-FLASH (1:10000) (Burch et al., 2011); mouse anti-phospho Mpm2 (source, 1:10000); rabbit anti-Slbp (Skrajna et al., 2017) (1:200); goat anti-rabbit Alexa 555 and 488 (Life Technologies, 1:400 and 1:200 respectively); goat anti-mouse Alexa 555 and 488 (Life Technologies, 1:400 and 1: 200 respectively).

### RNA and protein analysis

Histone mRNAs were analyzed by Northern blotting using probes to the coding region of either dH3 or dH1. As a loading control, a probe to the coding region of 7SK RNA was used, Five micrograms of total RNA was resolved on a 6% acrylamide-7M gel at 500V with 1X Tris/Borate/EDTA buffer (TBE). The samples contained xylene cyanol dye. After the dye had tun off the gel (∼1:10hr), the gel was run for another 30 minutes. The RNA was transferred to was transferred into a Positively Charged Nylon Transfer Membrane (GE Healthcare) by electroblotting. The membrane crosslinked with UV crosslinked in a Stratalinker (3X at (12000 units) and the hybridized with probes to histone mRNA. The RNA was detected by autoradiography or with a Phosphorimager. The filter was then hybridized to a probe for 7SK RNA. In some experiments the two probes were mixed prior to hybridization.

Proteins were resolved on an 8% polyacrylamide-SDS gel, transferred to nitrocellulose, incubated with the antibody to SLBP and the SLBP detected using ECL reagent. The membrane was then incubated with the β-actin antibody (GENETEX) as a loading control.

### *In situ* hybridization

Ovaries were processed for in situ hybridization using a published protocol (Goldman et al., 2019). Dissected ovaries were fixed with 4% formaldehyde for 20 mins. Next, the ovaries were washed with PBST and teased apart using a pipette. The ovaries were washed with 100% methanol for 5 min, then stored for 1 hour in 100% methanol at -20°C. The samples were then gradually re-hydrated into PBST. The samples were then washed for 10 minutes in Wash Buffer (4xSSC, 35% deionized formamide, 0.1% Tween-20). Probes were diluted in Hybridization Buffer (10% dextran sulfate, 0.1mg/ml salmon sperm ssDNA, 100 µl vanadyl ribonucleoside (NEB biolabs), 20ug/ml RNAse-free BSA, 4x SSC, 0.1% Tween-20, 35% deionized formamide) and were incubated with the sample overnight at 37°C. The following day, the samples were washed twice with Wash Buffer for 30 min. After two rinses with PBST and staining with DAPI, the ovaries were mounted on slides using Prolong Diamond (Life Technologies) and imaged.

The probes used were obtained from Stellaris and have been described previously (Hur, 2019).

### Immunofluorescence

Ovaries were processed for immunofluorescence using a previous described procedure (Goldman et al., 2019). Briefly, flies were fattened on yeast pellets for 3 days. Ovaries were fixed in 4% formaldehyde (Pierce) for 20 min. Primary antibody was incubated in 1× PBST (PBS + 0.1% Triton X-100) + 0.2% BSA (Promega) overnight at 4°C. The samples were then washed three times in PBST. The secondary antibody diluted in 1× PBST + 0.2% BSA was incubated with the sample overnight at 4°C. Following washes in PBST, the samples were mounted onto slides with Prolong Diamond (Life Technologies). For combined in situ and immunofluorescence, the ovaries were first processed for immunofluorescence. After removal of the secondary antibody, the samples were fixed in 4% formaldehyde for 5 min. Next, the samples were placed in 100% methanol for 1 hour at -20°C. After this incubation, the samples were processed for in situ hybridization as described in the previous section.

### Microscopy

Images were captured on a Zeiss LSM 780 inverted microscope equipped with Airyscan (Augusta University Cell Imaging Core). Images were processed for presentation using Adobe Photoshop, and Adobe Illustrator.

## ACKNOWLEDGMENTS

This work was supported by NIH grants R01GM58921 (W.F.M), R01 GM29832-41S1 (W.F.M. and J. P.-B. and R01GM100088 to G.G. J. P.-B. also received support from NIH grnat R25GM055336. We thank Stefano Di Talia (Duke Univ.), and Bob Duronio and Jim Kemp (UNC) for the in situ probes, and Xiao Yang for purification of the SLBP antibody.

