## supplemental material for "A region of Drosophila SLBP distinct from the histone pre-mRNA binding and processing domains is essential for deposition of histone mRNA in the oocyte"

SUPPLEMENTAL DATA

Supplemental Fig. 1. The $SLBP^{30}$ mutant embryos cannot complete the syncytial cell cycles. 0-2 hr embryos were collected from WT, $SLBP^{10}$ and $SLBP^{30}$ female flies and stained with DAPI. WT type embryos develop normally with a regular pattern of nuclei. As previously reported, embryos from $SLBP^{10}$ females fail to develop due to mitotic defects resulting from a low level of histone proteins (Sullivan et al., 2001). Embryos from $SLBP^{30}$ females have a similar phenotype but development arrests early than $SLBP^{10}$ embryos.

Supplemental Figure 2. (A-D) Early stage egg chambers from wild-type (A, B), $SLBP^{10}$ (C), and $SLBP^{\Delta 30}$ (D) were processed for in situ hybridization with the H3 coding region probe (red, H3-C). The mutant egg chambers were also processed for immunofluorescence using the FLASH antibody (green, C', D') and counterstained with DAPI (grey, C', D'). (E) A stage 10b wild-type egg chamber stained with antibodies against phosphor Mpm2 (red, E) and FLASH (green, E'). A merge image along with a DAPI (grey) counterstain is shown in E'' for a WT stage 10b egg chamber. This is an example of a stage 10b egg chamber that is positive for nuclear phospho Mpm2 as well as FLASH foci (arrows). 14% of the wild type egg chambers gave this pattern while 86% did not stain with mpm2.

Supplemental Figure 3. (A-D) Stage 10b egg chambers expressing a control shRNA (A, B) or an shRNA against *slbp* (C, D) were stained using the antibody against Slbp. The signal for Slbp is reduced upon expression of the *slbp* shRNA.

**Phosphorylation in the C-terminus of SLBP is not necessary for 3'end processing in vivo**

SLBP contains an RNA binding domain that is ~55% identical between Humans and Drosophila. However, the region C-terminal to the RBD, which also is required for processing, differs completely from the vertebrate SLBPs. The 17 aa after the RBD in Drosophila contains 4 serine's that are phosphorylated in vivo and necessary to properly process histone pre-mRNAs *in vitro* (Fig.S4C) (Dominski et al., 2002; Skrajna et al., 2017; Zhang et al., 2014). In vitro studies of point mutants either abolishing or mimicking phosphorylation of the four serines showed that the 4S-4A point mutant was inactive in histone pre-mRNA processing in vitro. Histone pre-mRNAs processing and the recruitment of the histone cleavage complex machinery in nuclear extracts depleted of SLBP, and the 4S-4E rescued this phenotype (Skrajna et al., 2017). The same study showed that the C-terminus of SLBP is not interchangeable, chimeras containing the Drosophila N-terminus and the RNA binding domain but the C-terminus of the Human SLBP abolishes processing in vitro (Skrajna et al., 2017).

To analyze the importance of these 4 Serine's in vivo we created point mutants to either mimic or abolish phosphorylation (Fig.S4C). Transgenes containing a $SLBP^{WT}$, the $SLBP^{4S\text{-}4E}$ or the $SLBP^{4S\text{-}4A}$ were integrated by recombinase mediated cassette exchange (RMCE) at the site 25C in the 2nd chromosome. In addition we mutated the conserved phosphorylation site in the TPNK in the RBD to NPNK. Genetic analyses were done to ask whether these transgenes could rescue the null mutant of SLBP, which dies in the first instar larvae. The wild type transgene rescued viability and fertility. Surprisingly both mutating the 4 serines to either A or to E in the C-terminus of SLBP, did not affect the ability of these SLBPs to rescue the null mutant. Ovaries were dissected from females expressing each transgene and used for western and northern blot analyses. We didn't observe any changes in protein levels for SLBP with each of the transgenes as shown in Fig.S4D. In addition neither point mutant showed production of polyadenylated histone mRNAs (Fig.S4E). In contrast the mutation of the TPNK to NPNK did not rescue the null mutant of SLBP, and the animals expressing this mutant transgene died in the first instar larva.  These results demonstrate that phosphorylation in the four serine's located in the C-terminus of SLBP are not required for histone pre-mRNA processing in vivo, although the mutations affected processing in vitro.

**Supplemental Fig. 4**.  Mutating phosphorylation sites in SLBP did not affect histone pre-mRNA metabolism.

**A**. We created mutations in the two known phosphorylation sites between aa 113 and 122 that are deleted in the SLBP30 mutant.  The transgenes were introduced at site 25C on chromosome 2.

**B.**  Both mutant flies developed normally and both the males and females were fertile. RNA was prepared from ovaries of adult female flies and analyzed by Northern blotting with the dH1 probe. No polyadenylated histone mRNA was detected.

**C.** Diagram of mutations in the C-terminus of SLBP.

**D.**  Western blots of SLBP from ovaries of WT flies or flies expressing only the transgenic SLBP from 12-16 hr embryos.  The lower molecular weight band (*)is a proteolytic product of SLBP. Actin was probed as a loading control.

**E.**  Total RNA was prepared from ovaries of flies expressing WT or the indicated mutant SLBPs. The blot was probed for histone dH3 mRNA.

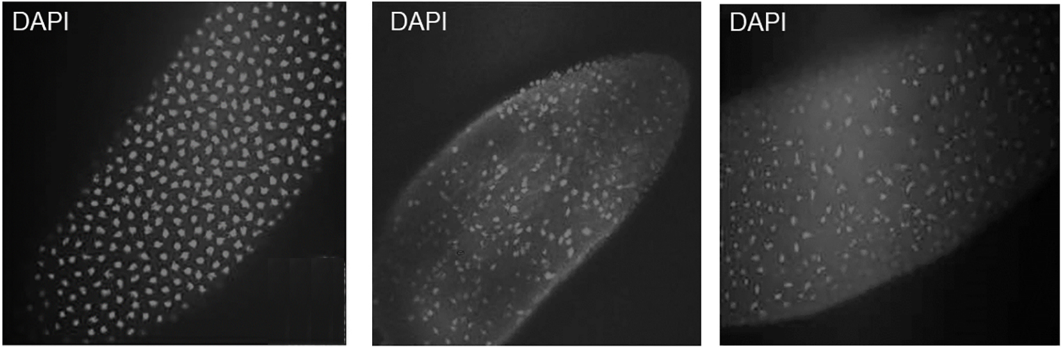


Wild type $SLBP^{10}$ $SLBP^{\Delta 30}$

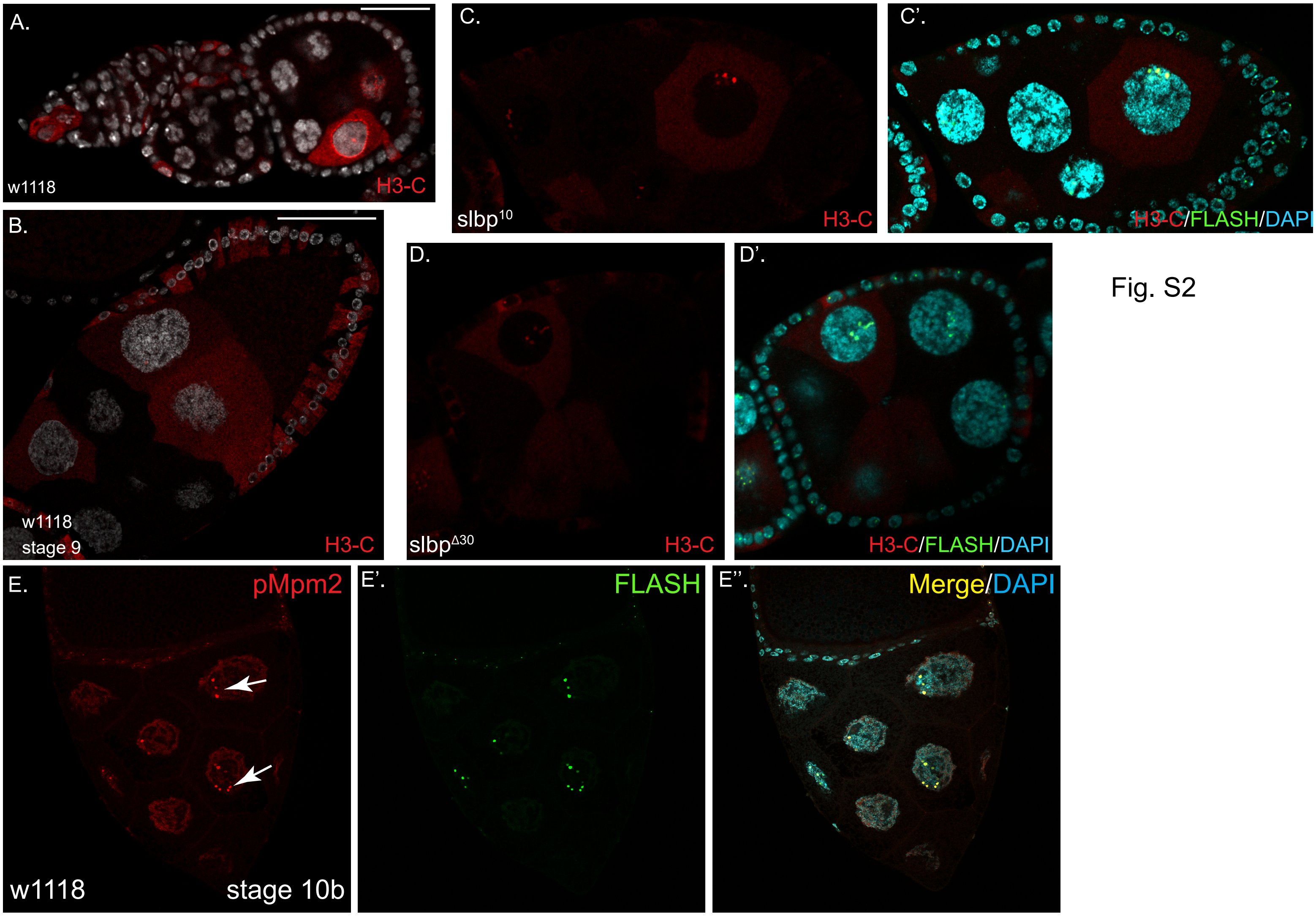


Fig. S2

Gonsalvez_Supplemental_Fig.1

Figure S3

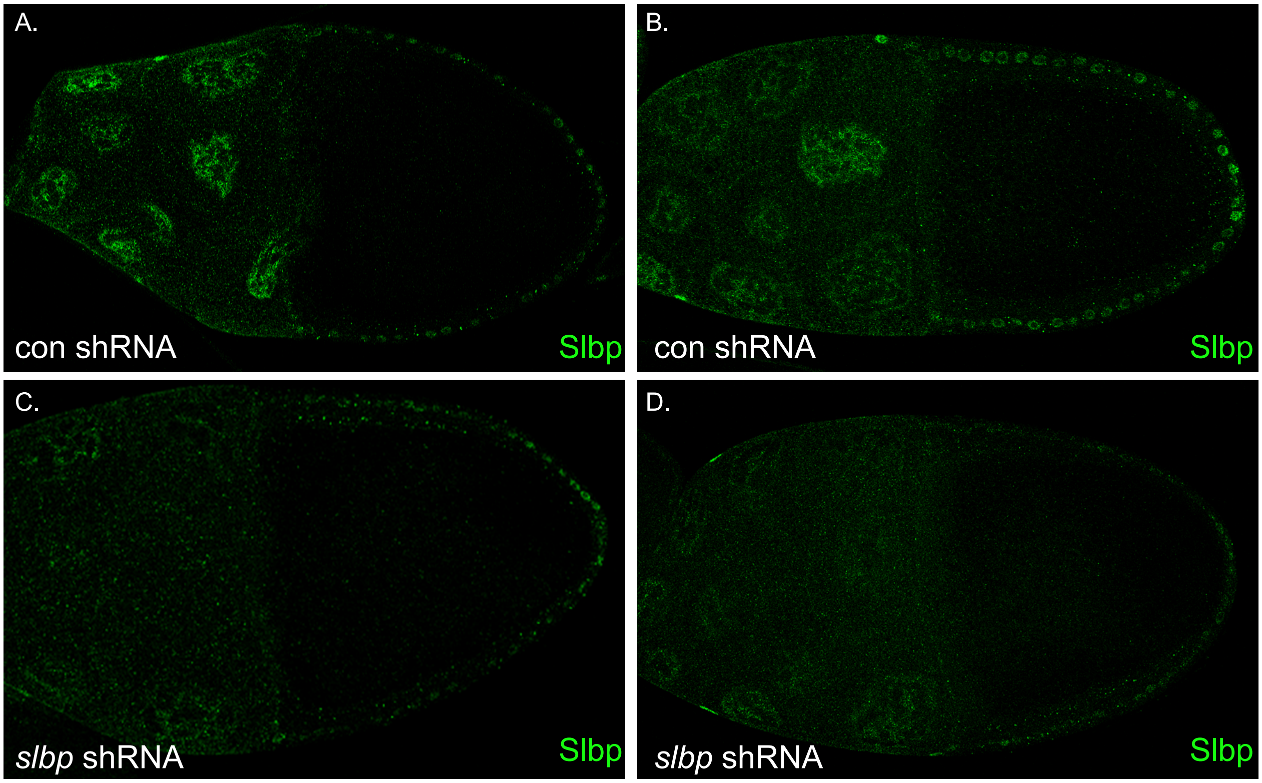


Gonsalvez_Supplemental_Fig.2

Fig. S4

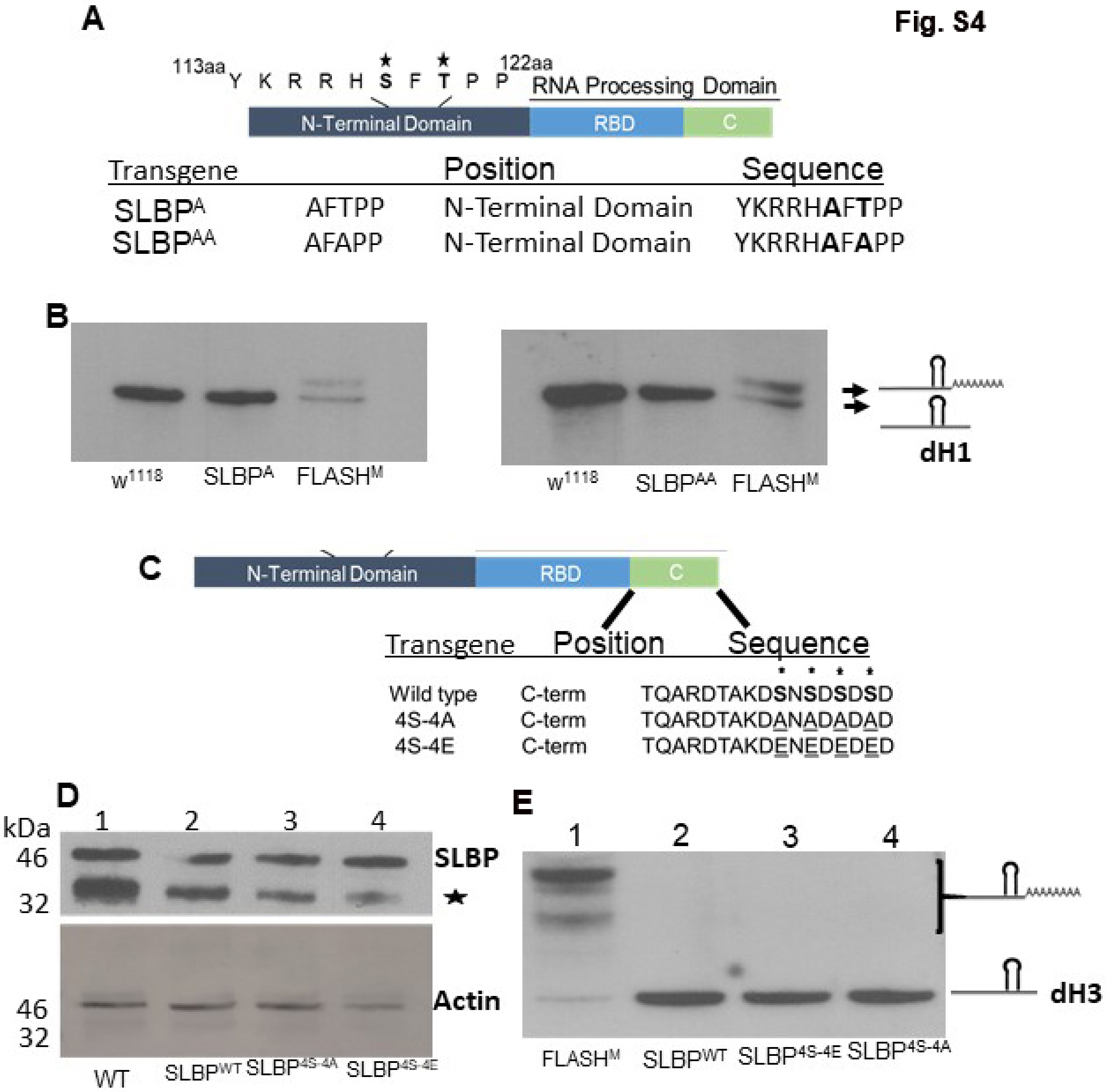
A
113aa Y K R R H S F T P P 122aa
RNA Processing Domain
N-Terminal Domain
RBD
C
Transgene
Position
Sequence
SLBP^A AFTPP N-Terminal Domain YKRRHAFTPP
SLBP^AA AFAPP N-Terminal Domain YKRRHAFAPP
B
w1118
SLBPA
FLASHM
w1118
SLBPAA
FLASHM
dH1
C
N-Terminal Domain
RBD
C
Transgene
Position
Sequence
Wild type C-term TQARDTAKDSNSDSDSD
4S-4A C-term TQARDTAKDANADADAD
4S-4E C-term TQARDTAKDENEDEDED
D
kDa
1
2
3
4
46
32
SLBP
Actin
46
32
WT
SLBPWT
SLBP4S-4A
SLBP4S-4E
E
1
2
3
4
dH3
FLASHM
SLBPWT
SLBP4S-4E
SLBP4S-4A
